# A single-cell map of antisense oligonucleotide activity in the brain

**DOI:** 10.1101/2023.02.14.528473

**Authors:** Meredith A Mortberg, Juliana E Gentile, Naeem Nadaf, Charles Vanderburg, Sean Simmons, Dan Dubinsky, Adam Slamin, Salome Maldonado, Caroline L Petersen, Nichole Jones, Holly B Kordasiewicz, Hien T Zhao, Sonia M Vallabh, Eric Vallabh Minikel

## Abstract

Antisense oligonucleotides (ASOs) dosed into cerebrospinal fluid (CSF) distribute broadly throughout the brain and hold the promise of treating myriad brain diseases by modulating RNA. CNS tissue is not routinely biopsied in living individuals, leading to reliance on CSF biomarkers to inform on drug target engagement. Animal models can link CSF biomarkers to brain parenchyma, but our understanding of how individual cells contribute to bulk tissue signal is limited. Here we employed single nucleus transcriptomics on tissue from mice treated with RNase H1 ASOs against *Prnp* and *Malat1* and macaques treated with an ASO against *PRNP*. Activity was observed in every cell type, though sometimes with substantial differences in magnitude. Single cell RNA count distributions implied target suppression in every single sequenced cell, rather than intense knockdown in only some cells. Duration of action up to 12 weeks post-dose differed across cell types, being shorter in microglia than in neurons. Suppression in neurons was generally similar to, or more robust than, the bulk tissue. In macaques, PrP in CSF was lowered 40% in conjunction with *PRNP* knockdown across all cell types including neurons, arguing that a CSF biomarker readout is likely to reflect disease-relevant cells in a neuronal disorder.

## INTRODUCTION

Antisense oligonucleotides (ASOs) can, in principle, modulate the expression of nearly any gene in the central nervous system (CNS), and hold the potential to treat myriad diseases^1^. Typically short (∼18-20 nucleotides) and chemically modified to improve pharmacokinetics and potency, ASOs are dosed directly into cerebrospinal fluid (CSF)^2^. Upon binding cell surface proteins, ASOs become internalized into endosomes, from which they escape gradually over weeks^3^ and achieve sustained pharmacologic activity in the cytoplasm and nucleus^4^. An ASO designed to modulate pre-mRNA splicing — nusinersen for spinal muscular atrophy — received FDA approval in 2016^5^. 23 ASOs have entered clinical trials for CNS disorders^1,6^, with several advancing to Phase III, along with others administered on an “N-of-1” basis^7,8^. The majority of CNS ASOs in trials today are “gapmers” — ASOs with 2 ‘ sugar modifications in the wings (typically 5 base pairs on either side) and a “gap” in the middle with no modifications except for a phosphorothioate backbone^2^ — designed to lower the expression of a target RNA by recruiting the enzyme RNase H1 to cleave it^9–11^. For CNS diseases caused by a toxic gain of function, gapmer ASOs offer a rational approach to target the root cause of disease by lowering the toxic RNA or protein^1^.

ASOs administered by bolus injection into CSF distribute broadly throughout the spinal cord and brain in rodents and monkeys^12,13^, albeit with a drug concentration gradient from superficial to deep brain structures. Animal models of CNS diseases with various regional and cell type-specific pathologies have been phenotypically ameliorated or modulated with ASO treatment^14–20^, and an ASO was shown active in at least four different cell types within mouse cortex at 2 weeks post-dose^12^. These findings suggest that ASO activity is relatively broadly distributed within the CNS. Our understanding of gapmer ASO activity across distinct cell types in the CNS remains limited, as the vast majority of target engagement data ever generated for ASOs are based on analyses of bulk tissue. This knowledge gap is particularly salient when considering the interpretation of CSF-based target engagement biomarkers in ASO trials. Any cell type-specific differences in ASO uptake or activity, combined with drug concentration gradients, could generate variability in the degree of target engagement among relevant CNS cells. And yet, a pharmacodynamic biomarker value in a single sampling compartment, such as target knockdown in CSF, may underpin choices to advance or halt clinical programs, and in pre-symptomatic prion disease, may even support regulatory determinations^21^. A deeper understanding of the profile of ASO activity across cell types should help to inform such crucial decisions.

Here, we employed single nucleus RNA sequencing (snRNA-seq) to characterize the distribution of ASO activity across individual cells and across cell types in the mouse and cynomolgus macaque CNS.

## MATERIALS AND METHODS

### Mice

All mice were female C57BL/6N. Animals for 3 week post-dose harvest were dosed at the Broad Institute (IACUC protocol 0162-05-17) and were 16 weeks old at the time of dosing. Animals for 2 and 12 week post-dose harvest were dosed at Ionis Pharmaceuticals (IACUC protocol 2021-1176) and were 8-12 weeks old at dosing. Mice were dosed via intracerebroventricular injection as described^22^. ASOs were delivered as a single bolus injection of 500 µg (*Prnp* ASOs) or 50 µg (*Malat1* ASO) formulated in a 10 µL volume of dPBS. Mice were perfused with HEPES-sucrose solution (110 mM NaCl, 10 mM HEPES, 25 mM glucose, 75 mM sucrose, 7.5 mM MgCl_2_, 2.5 mM KCl, pH 7.4) and brains harvested as described^23,24^.

### Non-human primates

Cynomolgus macaque (*Macaca fascicularis*) studies were performed at Labcorp Early Development Services GmbH (Münster, Germany) under IACUC protocol 8422120. Studies complied with all of the following regulations: European Directive 2001/83/EC, German Drug Law Arzneimittelgesetz, International Conference on Harmonization (ICH) gudelines M3(R2) (Guidance on Nonclinical Safety Studies for the Conduct of Human Clinical Trials and Marketing Authorization for Pharmaceuticals), ICH-S3A (Toxicokinetics: A Guidance for Assessing Systemic Exposure in Toxicology Studies), ICH-S4 (Duration of Chronic Toxicity Testing in Animals), and ICH-S8 (Immunotoxicity Studies for Human Pharmaceuticals). Animals were 2-4 years old at injection, mixed sex (2M/2F per cohort), and were of Asian origin. Lumbar punctures were performed on days 1, 29, 57, and 85. The procedure was performed fasting under ketamine/medetomidine anesthesia with a pencil-point pediatric needle at a position between L2 and L6. First, ≥0.5 mL of CSF was collected, then, 20 mg ASO was delivered in a 1 mL volume of artificial CSF (aCSF) injected over 1 minute, followed by a flush of 0.25 mL aCSF. 15 minutes after the procedure, animals were awakened with atipamezole. CSF was ejected into Protein LoBind tubes and snap frozen in liquid nitrogen. The CSF samples analyzed here were collected at day 85, just prior to the fourth dose, while brain tissues were collected at day 92. Because the majority of CSF volume was used for regulated studies, the aliquots available for analysis in this study varied from 120-300 µL and 0.03% CHAPS was added only after freeze/thaw; these pre-analytical factors likely contribute additional variability between samples^25^.

### Tissue dissection

For mouse brains, cryostat (Leica CM3050 S) dissection was performed as described^26^: after storage in optimal cutting temperature (O.C.T.) compound (Tissue-Tek 4583) at -80°C, mouse brains were mounted by the frontal cortex onto cryostat chucks with O.C.T. leaving the entire posterior half of the brain exposed. A ∼2.5 mg piece of tissue was then excised using a pre-chilled ophthalmic microscalpel (Feather P-715) and placed into a pre-chilled PCR tube. For mice, a piece of somatosensory cortex was used for snRNA-seq, while an adjacent piece of visual cortex was used for bulk qPCR; thalamus was cut along the fiber tract and the dorsal half was used for snRNA-seq while the ventral half was used for qPCR; cerebellum was cut through the ansiform lobule and a piece of simple/ansiform lobule was used for snRNA-seq while a piece of ansiform/paramedian lobule was used for qPCR. Cynomolgus brains were coronally sectioned at a thickness of 4 mm, and cylindrical tissue punches of 2 mm diameter were taken for RNA analysis and of 6 mm diameter for protein analysis. The 2 mm diameter by 4 mm length cylindrical tissue punch was then sectioned lengthwise into quarters on the cryostat and one quarter was used for single cell analysis. From the most rostral section containing frontal cortex, punches were taken from middle frontal gyrus across all histological cortical layers. From section 12, where both cerebellum and medulla are visible, punches were taken from the posterior lobe of the cerebellum across the granular, ganglionic and molecular layers.

### Bulk tissue qPCR

Tissue pieces dissected on the cryostat were placed in RNAlater-ICE (Invitrogen AM7030) and allowed to thaw overnight at -20°C. Once samples were thawed, tissue was homogenized in 1 mL QIAzol lysis reagent, using 3 × 40 second pulses on a Bertin MiniLys homogenizer in 7 mL tubes pre-loaded with zirconium oxide beads (Precellys CK14, Bertin KT039611307.7 / P000940-LYSK0-A). RNA was isolated from homogenate using RNeasy Lipid Tissue Mini Kit (Qiagen 74804) per the manufacturer protocol. RNA was eluted with 40 µL RNase-free water. RT-PCR samples were prepared using Taqman 1-Step RT-PCR master mix (Invitrogen) and Taqman gene expression assays (Invitrogen) for mouse *Prnp* (Mm00448389_m1; spanning exons 1-2) and mouse *Tbp* (Mm00446971_m1) and for cynomolgus *TBP* (Mf04357804_m1). The following gene-specific primer-probe sets were custom ordered from IDT: *Malat1* (mouse), Forward: AGGCGGGCAGCTAAGGA, Reverse: CCCCACTGTAGCATCACATCA, Probe: TTCCTCTGCCGGTCCCTCGAAAG; *PRNP* (cynomolgus; spanning intron 1 - exon 2), Forward: CCTCTCCTCACGACCGA, Reverse: CCCAGTGTTCCATCCTCCA, Probe: CCACAAAGAGAACCAGCATCCAGCA. Samples were run on a QuantStudio 7 Flex system (Applied Biosystems) using manufacturer ‘s recommended cycling conditions. Each biological sample was run in duplicate and the level of all targets were determined by ΔΔCt whereby results were first normalized to the housekeeping gene *Tbp* and then to PBS- or aCSF-treated animals.

### Single cell sequencing

After cryostat dissection, samples were batched in groups of eight, chosen to include treated and control animals in every run. Single nucleus suspensions were prepared as described^27,28^. Briefly: tissue samples were triturated, by pipetting, in an extraction buffer containing Kollidon VA64, Triton X-100, bovine serum albumin, and RNase inhibitor, then passed through a 26-gauge needle, washed and pelleted, then passed through a cell strainer. Nuclei positive for DAPI signal were isolated by fluorescence-activated cell sorting with a Sony SH800 or MA900 calibrated with a 70 µm chip, with a 405 nm excitation laser and light collected with a 425 - 475 nm filter. Sorted nuclei were counted using a Fuchs-Rosenthal C-Chip hemocytometer and a hand tally counter. A volume chosen to target 17,000 nuclei was submitted to the Broad Institute ‘s Genomics Platform, where 10X library construction (3 ‘ V3.1 NextGEM with Dual Indexing) was performed according to manufacturer instructions^29^. Libraries were sequenced on an Illumina Novaseq 6000 S2 for 100 cycles.

### Data processing and analysis

Raw binary base call (BCL) files were synced to Google Cloud and analyzed on Terra.bio. Cumulus^30^ Cell Ranger^31^ 6.0.1 (cellranger_workflow v28) was employed, with flags --include_introns and --secondary set to true, to process BCL files into unique molecular identifier (UMI) count matrices for each individual sample. Mouse samples were aligned to Cell Ranger reference package mm10-2020-A and cynomolgus samples were aligned to a custom Cell Ranger reference made from Ensembl *Macaca fascicularis* 6.0 (release 108). Matrices were then aggregated using Cell Ranger 7.0.1 (aggr with the --normalize flag set to none) to yield one UMI count matrix per species and brain region. Statistical analyses and data visualization were conducted using custom scripts in R 4.2.0.

### Cell type assignment

Aggregated count matrices were examined using Loupe Browser. Viewing cells in 2-dimensional uniform manifold approximation and projection (UMAP)^32^ space, we looked for cell type markers established or validated in several prior single-cell studies^26,33–39^. Clusters corresponding to empty droplets, doublets, debris, or mitochondria were flagged and removed based on low UMI or unique gene count, low percentage intronic reads, lack of obvious differentially expressed genes, high expression of mitochondrial genes, or location between two other clusters and expression of markers of each. Assignments were then validated by generating dot plots in Seurat V4^40^ in R. For cortical excitatory and inhibitory neurons in 3 week post-dose animals, a list of barcodes was exported from R and reclustered in Loupe Browser.

### Statistics

For each combination of brain region, timepoint, and treatment condition, snRNA-seq data were grouped by animal and cell type and the sum of target UMIs and total UMIs was calculated. A negative binomial model was fit to the resulting data, with target RNA UMIs as the dependent variable; cell type and a cell type-treatment interaction term as the dependent variables, and total UMIs as the offset. This utilized the MASS package in R, with the call: glm.nb(target_umi ∼ celltype + celltype:treatment + offset(log(total_umi))). This returns coefficients in natural logarithm space. For the ASO-treated conditions, the coefficient for each cell type-treatment interaction term coefficient was then exponentiated to yield the mean estimate of the residual target RNA in that cell type. The 95% confidence interval was defined as that mean estimate ±1.96 of the standard errors returned by the model. Each individual animal ‘s point estimate of residual target RNA in each cell type was obtained by adding the residual from the model to the cell type-treatment coefficient, and then exponentiating. To account for the different abundance of different cell types, which impacts the size of our confidence intervals on target knockdown, we used weighted Pearson ‘s correlations (wtd.cor from the weights package in R) to test candidate variables and weighted standard deviations (square root of wtd.var from the Hmisc package in R) to evaluate the variability in target engagement between cell types within different brain regions. Throughout, all error bars and shaded areas in figures represent 95% confidence intervals. P values less than 0.05 were considered nominally significant.

### Data availability

A public git repository will be made available at https://github.com/ericminikel/scaso containing all source code and a minimum analytical dataset (∼200 MB) sufficient to reproduce most figures and statistics in this manuscript. The full dataset (∼8 TB) will be deposited at singlecell.broadinstitute.org and made available through a public Terra repository.

## RESULTS

### Generation and cell type classification of single nucleus transcriptomes

We selected 4 previously characterized ASOs (Table 1) including 2 *Prnp* ASOs for which we have evaluated survival benefit in prion-infected mice^22,41^, 1 *Malat1* ASO for which an extensive regional pharmacokinetic and pharmacodynamic atlas has been published^12^, and 1 human *PRNP* ASO sequence-matched in cynomolgus macaques^42^. We analyzed a total of 78 single nucleus transcriptomes from tissues of mice and macaques treated with these ASOs or with vehicle (Table 2). For mouse *Prnp* tool compounds, we used a dose of 500 µg evaluated in survival studies^22,41^, whereas for the more potent *Malat1* ASO, we used a 50 µg dose^12^. Each sample yielded an average of 532 million reads mapping to 7,667 cells and yielding 7,650 unique molecular identifiers (UMIs) per cell, corresponding to a median of 3,108 detected genes per cell (Table S1).

**Table 1.** Compounds used in this study. Color code for ASO chemical modifications: black = unmodified deoxyribose (2′H; DNA). orange = 2′ methoxyethyl (MOE). blue = 2′-4′ constrained ethyl (cET). Unmarked backbone linkages = phosphorothioate (PS); linkages marked with o = normal phosphodiester (PO). mC = 5-methylcytosine.

| ASO | sequence and chemistry | target | ref |
| --- | --- | --- | --- |
| ASO 1 | mCToAoTTTAATGTmCAoGoTmCT | mouse *Prnp* 3'UTR | 22,41 |
| ASO 6 | mCToTomCoTATTTAATGTmCAoGoTmCT | mouse *Prnp* 3' UTR | 22 |
| *Malat1* ASO | GCoCoAoGoGCTGGTTATGAoCoTCA | mouse/NHP *Malat1* | 12 |
| ASO N | GToCoAoToAoATTTTCTTAGCoTAC | human/NHP *PRNP* intron | 42 |

**Table 2.** Summary of samples analyzed by single-cell sequencing. Identities of active compounds are shown in Table 1; aCSF = artificial CSF; PBS = phosphate-buffered saline. Note that weeks post-dose is after a single dose for mice, whereas cynomolgus macaques received 4 doses at 4 week intervals and were sacrificed after 13 weeks, 1 week after the final dose.

| species | treatment | dose | brain region | weeks post-dose | N animals |
| --- | --- | --- | --- | --- | --- |
| mouse | PBS | — | cortex | 2 | 4 |
| mouse | ASO 1 | 500 µg 1x | cortex | 2 | 4 |
| mouse | ASO 6 | 500 µg 1x | cortex | 2 | 4 |
| mouse | ASO 6 | 500 µg 1x | cortex | 3 | 4 |
| mouse | PBS | — | cortex | 3 | 7 |
| mouse | ASO 6 | 500 µg 1x | thalamus | 3 | 3 |
| mouse | PBS | — | thalamus | 3 | 4 |
| mouse | PBS | — | cerebellum | 3 | 4 |
| mouse | ASO 6 | 500 µg 1x | cerebellum | 3 | 4 |
| mouse | PBS | — | cortex | 12 | 4 |
| mouse | ASO 1 | 500 µg 1x | cortex | 12 | 4 |
| mouse | ASO 6 | 500 µg 1x | cortex | 12 | 4 |
| mouse | PBS | — | cerebellum | 12 | 4 |
| mouse | ASO 6 | 500 µg 1x | cerebellum | 12 | 4 |
| mouse | Malat1 ASO | 50 µg 1x | cerebellum | 12 | 4 |
| cynomolgus | aCSF | — | cerebellum | 13 | 4 |
| cynomolgus | ASO N | 20 mg 4x | cerebellum | 13 | 4 |
| cynomolgus | aCSF | — | cortex | 13 | 4 |
| cynomolgus | ASO N | 20 mg 4x | cortex | 13 | 4 |

Single nucleus transcriptomes were aggregated by species and brain region to yield five count matrices. These were projected into a two-dimensional space using uniform manifold approximation and projection (UMAP; Figure 1) revealing distinct clusters corresponding to specific cell types. Cell types assigned using established markers were validated using dot plots to visualize the specificity of gene expression (Figure 1).

**Figure 1.**
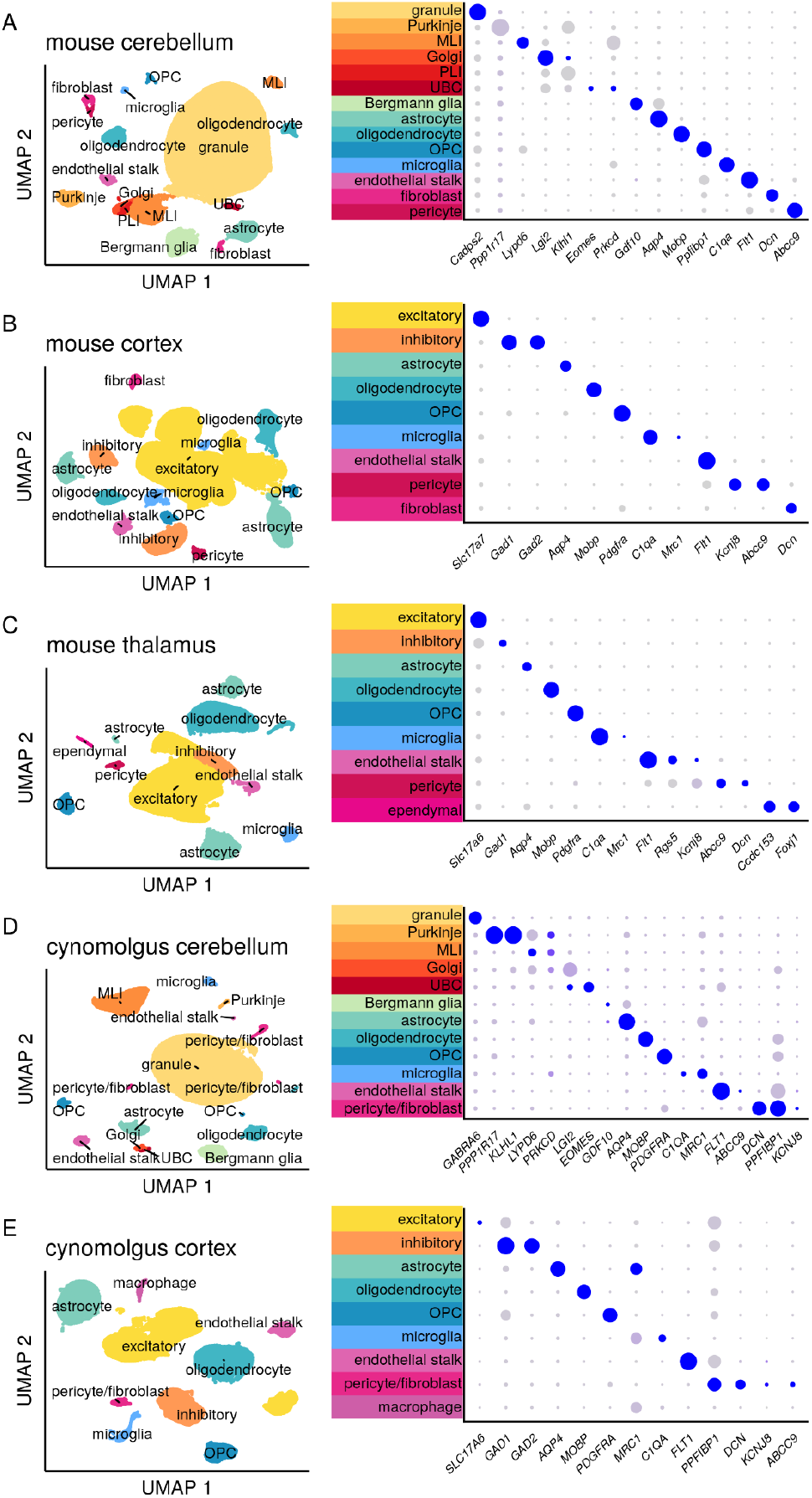
Clustering and assignment of brain cell types. Single-cell gene expression profiles projected into two dimensions using Uniform Manifold Approximation and Projection^32^ (UMAP) and characterized using dot plots. In dot plots, gray to blue color gradient represents higher expression while small to large dot size gradient represents broader expression. Thus, a large blue dot indicates a marker widely and highly expressed by cells within the indicated cluster; a small gray dot indicates little to no expression by those cells. A small blue dot can indicate a marker highly expressed by only a subset of cells within the cluster, while a large gray dot can indicate a marker broadly but lowly expressed. MLI = molecular layer interneuron, UBC = unipolar brush cell, PLI = Purkinje layer interneuron, OPC = oligodendrocyte progenitor cell. For breakdown by weeks post-dose and active/inactive treatment group see Figure S1.

### Distribution of ASO activity at the single cell level

The long non-coding RNA *Malat1* is a valuable model target for single cell assessment of ASO activity because it is highly expressed, accounting for 11.4% of all UMIs in our mouse transcriptomes, and because a potent and well-characterized tool ASO against *Malat1* is available^12^. Considering this compound ‘s median effective dose (ED_50_) of ∼50 µg in cerebellum^12^ and the prior evidence for some difference in activity between cerebellar cell types^12,19^, we examined *Malat1* knockdown in cerebellum at 12 weeks after a single 50 µg ICV dose of *Malat1* ASO. Aggregation of single nucleus sequencing data across all mouse cerebellar nuclei indicated 45.4% residual *Malat1*, close to the 52.4% residual detected in an adjacent piece of cerebellar tissue analyzed by bulk qPCR (Figure 2A; Table S2-S3). When *Malat1* UMIs per cell were visualized as a histogram (Figure 2B), the median cell possessed 360 *Malat1* UMIs in PBS-treated animals. ASO treatment yielded a bimodal histogram, with a main peak at 166 UMIs but a second peak at ∼35 UMIs (black arrow, Figure 2B) indicating deeper knockdown in a subpopulation of cells. Across 14 different cell types (Figure 2C), residual *Malat1* ranged from 7% to 76%, while each individual histogram appeared unimodal. This suggested that the bimodality in the histogram for bulk tissue (Figure 2B) is due to differences in knockdown in different cell types. Scatterplots of *Malat1* UMIs vs. total UMIs indicated that the width of the distributions in these histograms is largely due to differences in total UMIs per cell. All of these observations were consistent with broad knockdown in every detected cell. Fitting residual *Malat1* in each cell type in each sample with a negative binomial model (see Materials and Methods) confirmed substantial differences in knockdown between cell types (Figure 2D; Table S4-S6).

**Figure 2.**
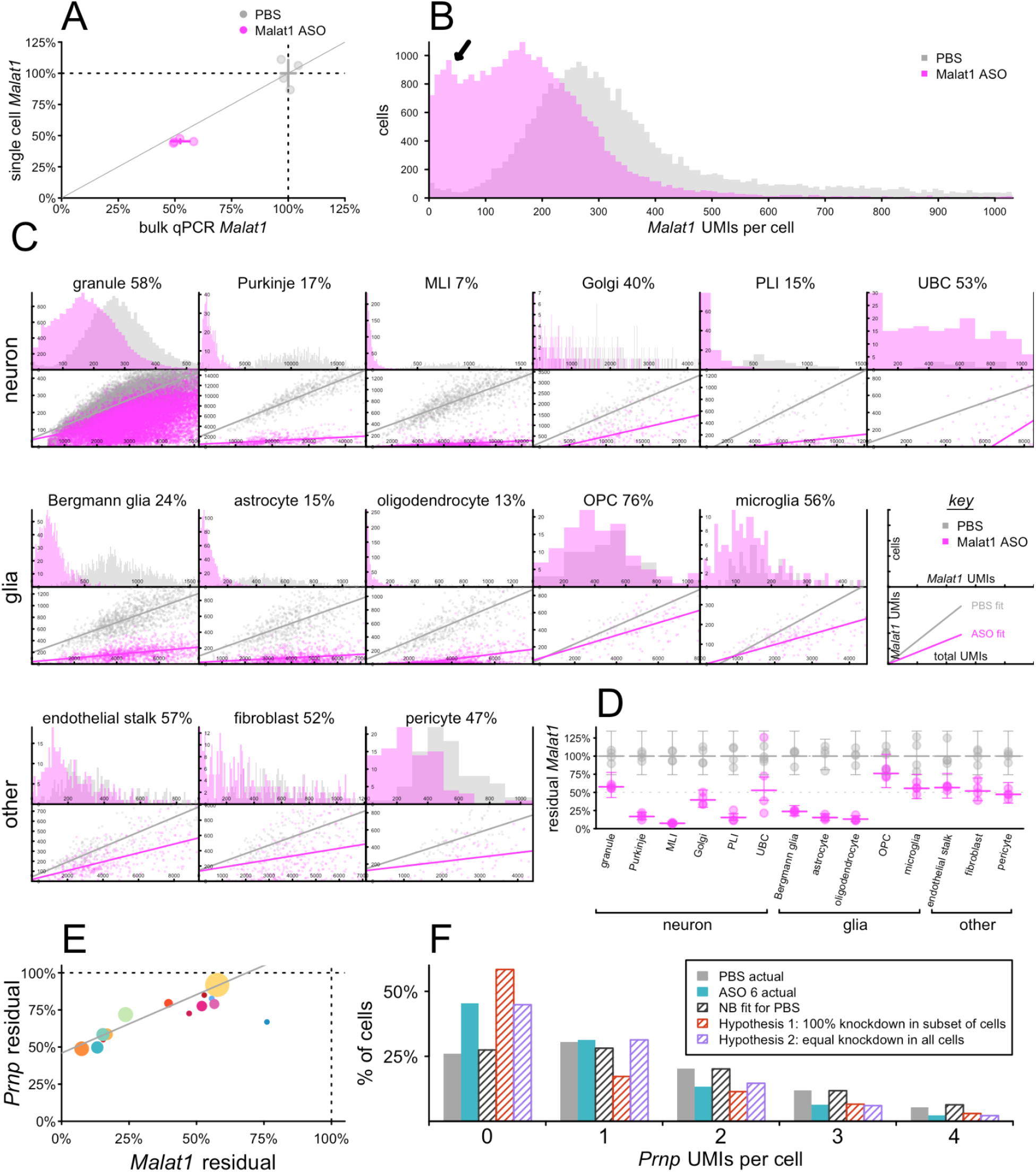
Single-cell distribution of ASO activity in mouse cerebellum at 12 weeks post-dose. N = 4 mice per group received 50 µg Malat1 ASO, 500 µg ASO 6, or PBS ICV and cerebella were harvested 12 weeks later. **A)** Malat1 knockdown after Malat1 ASO treatment, assessed by bulk qPCR (x axis) versus aggregation of single cell sequencing regardless of cell type (y axis), for individual animals (points) and groups (crosshairs indicate means and 95% confidence intervals). **B)** Histogram of the number of Malat1 UMIs per single cell across all samples. Arrow indicates a second peak observed only for treated animals. **C)** Breakdown across 14 cell types. Top panels are histograms of single cells as in B, but broken down by cell type. Bottom panels are scatterplots showing total UMIs per single cell versus Malat1 UMIs per cell and best fits by linear regression (Table S8). Percentages indicate residual Malat1. Note the “key” panels at the far right of the middle row. **D)** Negative binomial (NB) modeling of single cell data as point estimates of knockdown for each animal and each cell type (points) and means and 95% confidence intervals (bars) for each treatment group and cell type. **E)** Correlation across cell types of Prnp knockdown by ASO 6 and Malat1 knockdown by Malat1 ASO. Each point is a cell type, colors are from Figure 1A, and point sizes are logarithmically scaled with number of cells sequenced. **F)** Histogram of Prnp UMIs per single cell in astrocytes for PBS (gray) and ASO 6-treated animals (cyan). Shaded gray bars indicate the distribution predicted by a NB model fit to the PBS data. Shaded red bars indicate the distribution if the observed 56% residual Prnp RNA in astrocytes corresponded to 56% of cells following the original NB distribution and 44% being set to zero. Shaded cyan bars indicate the distribution if the observed 56% residual corresponded to the NB parameter mu being reduced by 44%.

snRNA-seq inherently yields low sequencing coverage in any one nucleus: our count matrices, measuring ∼30,000 genes long by ∼60,000 - 270,000 nuclei wide, were 87.2 - 93.2% populated by zeroes (Table S7). In other words, most genes are not detected in most nuclei, even where they are expressed. Thus, unlike *Malat1*, whose expression in single nuclei was normally distributed, most potential ASO targets will have UMI counts that are Poisson or negative binomial distributed in single nuclei data. For instance, *Prnp* averaged just 0.85 UMIs/cell in the cerebella of PBS-treated animals. Nevertheless, when the data from 12 weeks after a single 500 µg dose of ASO 6, were fit to the same negative binomial model as *Malat1*, ASO 6 displayed a highly similar pattern of activity across cerebellar cell types (rho = 0.96, P < 3.9e-8, weighted Pearson ‘s correlation; Figure 2E; Table S6).

Despite the lower basal expression of *Prnp*, we posited that examination of the histograms of UMIs/cell for *Prnp* could reveal information about the distribution of drug activity across single cells. As an example, we considered astrocytes, which had 56% residual *Prnp*, comparing histograms of actual *Prnp* UMIs/cell in PBS and ASO 6-treated animals versus three models. A negative binomial model fit to the PBS-treated animals mirrored these animals ‘ actual distribution almost perfectly. Lowering *Prnp* to 56% residual by setting 44% of astrocytes ‘ *Prnp* counts to zero would have yielded a histogram with far more zeroes, and fewer ones, than the observed distribution in ASO 6-treated animals. In contrast, lowering *Prnp* to 56% residual by lowering the negative binomial parameter mu by 44%, corresponding to equal knockdown in all cells, yielded a distribution that closely mirrored the observed distribution in ASO-treated animals (Figure 2F; Table S9). Thus, for *Prnp* as for *Malat1*, bulk tissue knockdown appears to arise from broad knockdown in all cells, albeit with a stereotypical pattern of differences across distinct cell types.

### ASO activity across regions and cell types in the mouse brain

We assessed the profile of ASO target engagement across cell types in 3 brain regions in mice at 3 weeks post-dose with *Prnp* ASO 6 (Figure 3). Because this tool compound is less potent than the *Malat1* ASO, we used a 500 µg dose, which modifies prion disease in mice^22,41^ and lowers whole hemisphere PrP to an estimated 56% residual after 4 weeks^43^. Whereas *Malat1* localizes to the nucleus^44^, *Prnp* is a protein-coding gene whose mRNA reaches the cytosol, and ASO 6 targets the *Prnp* 3 ‘UTR, so cytosolic activity is possible. Nonetheless, we found that the value of residual *Prnp* obtained by snRNA-seq, which will detect nuclear ASO activity only, agreed closely with the value obtained by bulk tissue qPCR, which used exon junction-spanning primers and therefore will only detect mature mRNA (Figure 3A-C).

**Figure 3.**
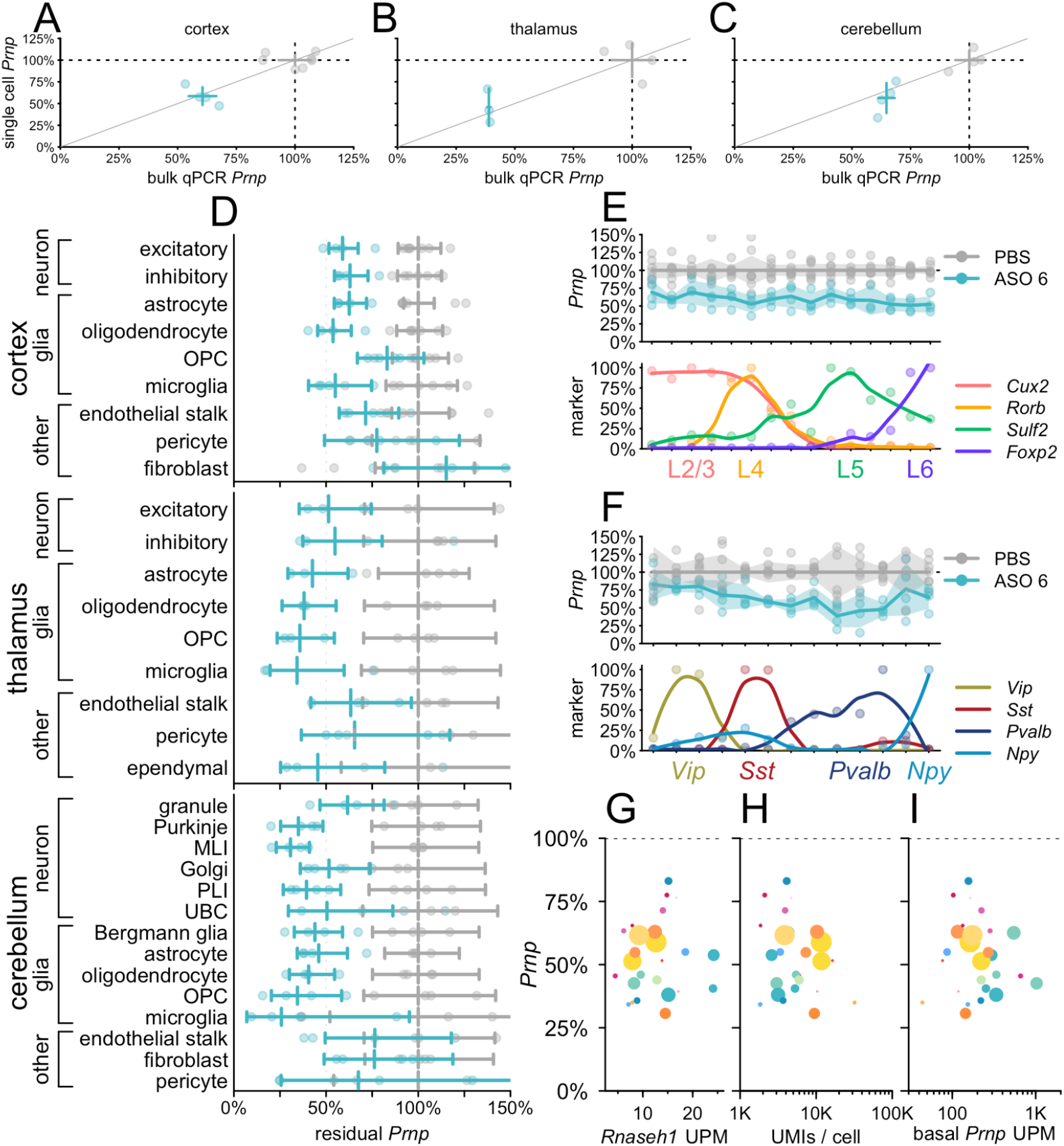
Atlas of ASO activity across cell types in mouse brain at 3 weeks post-dose. Mice 3 weeks after a single 500 µg dose of ASO 6. **A-C)** Concordance of bulk qPCR (x axis) and single cell (y axis) measurement of total Prnp knockdown in cortex (A), thalamus (B), and cerebellum (C). Points are individual animals, crosshairs are 95% confidence intervals on both dimensions. **D)** Residual Prnp expression across cell types in three brain regions. Error bars are 95% confidence intervals of the mean. **E)** Cortical excitatory neurons from panel A were reclustered into 15 clusters ranked by expression of four excitatory layer markers. Residual Prnp (top) is shown as individual animals (points), means (lines), and 95% confidence intervals of the mean (shaded areas). Each marker ‘s expression (bottom) is normalized to the cluster with the highest expression; points are normalized values and curves are loess fits. **F)** Cortical inhibitory neurons from panel A, reclustered and plotted as in panel B. **G-I)** Scatterplots of residual Prnp expression (y axis) versus candidate covariates (x axes). Each point represents a region and cell type combination from panel D (cortical fibroblasts, the only cell type with nominally >100% residual expression, are not visible). Colors correspond to cell type colors in Figure 1, and dot sizes scale logarithmically with the number of cells sequenced. Candidate covariates examined are Rnaseh1 expression in UMIs per million (UPM; G), total UMIs per cell (H), and basal Prnp expression (UPM; I).

Breakdown of single cell data by cell type showed broad target engagement across cell types including diverse types of neurons and glia (Figure 3D). As with the *Malat1* ASO (Figure 2), cell type differences were relatively pronounced in the cerebellar neurons, where knockdown was deeper in Purkinje cells and MLI than in granule cells. Across regions in ASO-treated animals, endothelial stalk cells, pericytes, and fibroblasts generally had both the highest residual *Prnp* and the lowest count of cells sequenced, giving rise to wide confidence intervals that overlapped the PBS-treated animals. Nonetheless, point estimates for these cells generally suggested some target engagement, with the possible exception of cortical fibroblasts. To further examine the profile of knockdown among neuronal subtypes, we reclustered cortical excitatory (Figure 3E; Table S10) and inhibitory (Figure 3F; Table S11) neurons and ordered them by relative expression of excitatory layer or inhibitory subtype markers. Target engagement appeared similar across all layers of excitatory neurons (Figure 3E). Knockdown appeared possibly deeper in *Pvalb*-expressing than in *Vip*-expressing inhibitory neurons, but again, target engagement was observed across all subtypes (Figure 3F).

Across all regions, we asked whether the differences in residual *Prnp* across cell types could be explained by any obvious candidate variables. *Rnaseh1* expression varied little among cell types and did not predict knockdown (P = 0.61, weighted Pearson ‘s correlation; Figure 3G).

Total UMIs per cell, a potential proxy for cell size^26^, and basal *Prnp* expression, were likewise uncorrelated (P = 0.64 and P=0.08, weighted Pearson ‘s correlation; Figure 3H-I; see Discussion).

### Potency and duration of action across ASO chemistries

Gapmer ASOs currently in clinical trials are 2 ‘MOE gapmers similar to ASO 6, but improved chemical modifications of ASOs are a highly active area of research^45,46^, prompting us to investigate the cell type profile of an ASO incorporating 2 ‘-4 ‘ constrained ethyl (cEt) modifications^47^. *Prnp* ASO 1 (Table 1), a mixed 2 ‘MOE/cEt oligonucleotide, targets the same site as ASO 6 and is effective in prion-infected mice^22,41^. We evaluated the activity of ASO 1 and ASO 6 in mouse cortex at both 2 and 12 weeks after a single 500 µg bolus dose (Figure 4). As before, despite the 3 ‘UTR target of these ASOs, single cell and bulk qPCR measurements of overall knockdown were in reasonable agreement (Figure 4A). ASO 1 had a shorter duration of action than ASO 6, with residual target rising from 47% to 91% of saline controls (by bulk qPCR) residual, a 44% recovery, versus 31% to 65%, a 34% recovery, for ASO 6 (Figure 4A; Table S3). Each compound provided substantial knockdown at 2 weeks across all cell types detected, and each exhibited marked differences across cell types in the rate of recovery (Figure 4B; Table S12). For example, for both ASOs, microglia exhibited the most complete recovery of any cell type (+51% for ASO 6 and +70% for ASO 1), while excitatory neurons were comparatively steady (+29% for both; Figure 4B).

**Figure 4.**
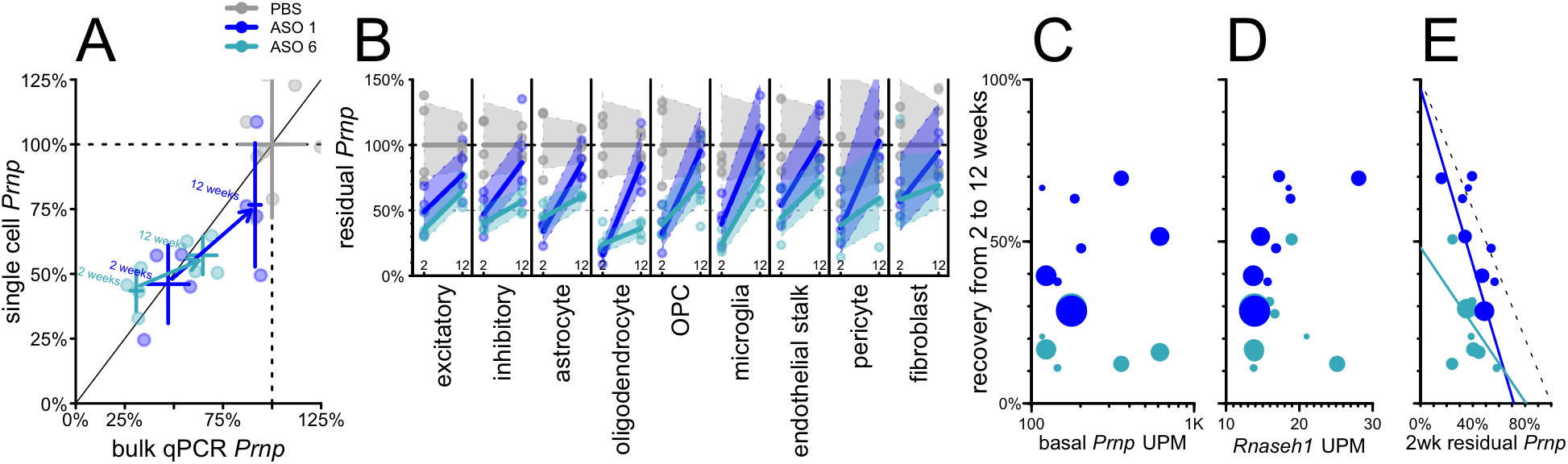
Duration of activity across different ASO chemistries in mouse cortex from 2 to 12 weeks post-dose. Groups of N = 4 mice received PBS, 500 µg ASO 1, or 500 µg ASO 6 and cortex was evaluated after 2 or 12 weeks. **A)** Overall residual Prnp assessed by bulk qPCR (x axis) versus aggregate single cell data without regard to cell type (y axis). Points represent individual animals and crosshairs represent means and 95% confidence intervals on both axes. 2 week data (lower left) and 12 week data (upper right) are connected by arrows indicating washout for both ASO 1 (dark blue) and ASO 6 (cyan). **B)** Washout from 2 weeks (left) to 12 weeks (right) for each cell type in mouse cortex for ASO 1 and ASO 6. Lines represent means and shaded areas represent 95% confidence intervals. **C-E)** Scatterplots of candidate variables (x axis) basal Prnp expression (UMIs per million, UPM; C), Rnaseh1 expression (UPM; D), and 2 week residual Prnp (E) versus percentage points of recovery (washout at 12 vs. 2 weeks, y axis) for each cell type (cell types are points, sized logarithmically by number of cells) for both ASO 1 (dark blue) and ASO 6 (cyan).

Across both compounds, neither basal *Prnp* expression (RPM in PBS-treated animals) nor *Rnaseh1* expression showed any correlation with washout between 2 and 12 weeks (P = 0.78 and P = 0.12, weighted Pearson ‘s correlation; Figure 4C-D). The depth of target suppression at 2 weeks post-dose, however, showed an inverse correlation with recovery by 12 weeks which was significant for ASO 1 (rho = -0.84, P = 0.0043, weighted Pearson ‘s correlation) and directionally consistent for ASO 6 (rho = -0.41, P = 0.27, weighted Pearson ‘s correlation; Figure 4E; see Discussion).

### Cell type profile and biomarker impact in non-human primates

We examined tissue from cynomolgus macaques that received ASO N. In addition to permitting us to examine ASO activity in a larger brain, the macaques also differed from our mice in being dosed intrathecally (IT) rather than ICV, and receiving 4 repeat doses at 4-week intervals. In cortex, bulk residual *PRNP* measured by snRNA-seq again mirrored that by qPCR (Figure 5A), although in cerebellum, knockdown measured by snRNA-seq appeared slightly deeper (Figure 5B). Residual PrP protein level quantified by ELISA^43^ in ASO N-treated animals was 41% in cortex, 82% in cerebellum and 60% in CSF (Figure 5C; Table S13-S14). Target engagement was broadly observed across all detected cell types in both cortex and cerebellum (Figure 5D; Table S6). In cortex, knockdown was deepest in neurons and weakest in endothelial stalk and pericytes/fibroblasts. In cerebellum, knockdown was deepest in Purkinje cells and molecular layer interneurons (MLIs) and weakest in pericytes/fibroblasts. Because these tissues were obtained just 1 week after the animals ‘ final dose of ASO, we compared the cell type profile of target engagement in macaques to that observed in mice 2 weeks after a single dose of ASO 6 (Figure 5E-F). The two datasets shared robust knockdown in the aforementioned neuronal populations and relatively limited knockdown in pericytes/fibroblasts. Correlation of residual target percentages across cell types was positive but non-significant for cortex (rho = 0.40, P = 0.33, weighted Pearson ‘s) and positive and significant for cerebellum (rho = 0.80, P = 0.0019, weighted Pearson ‘s).

**Figure 5.**
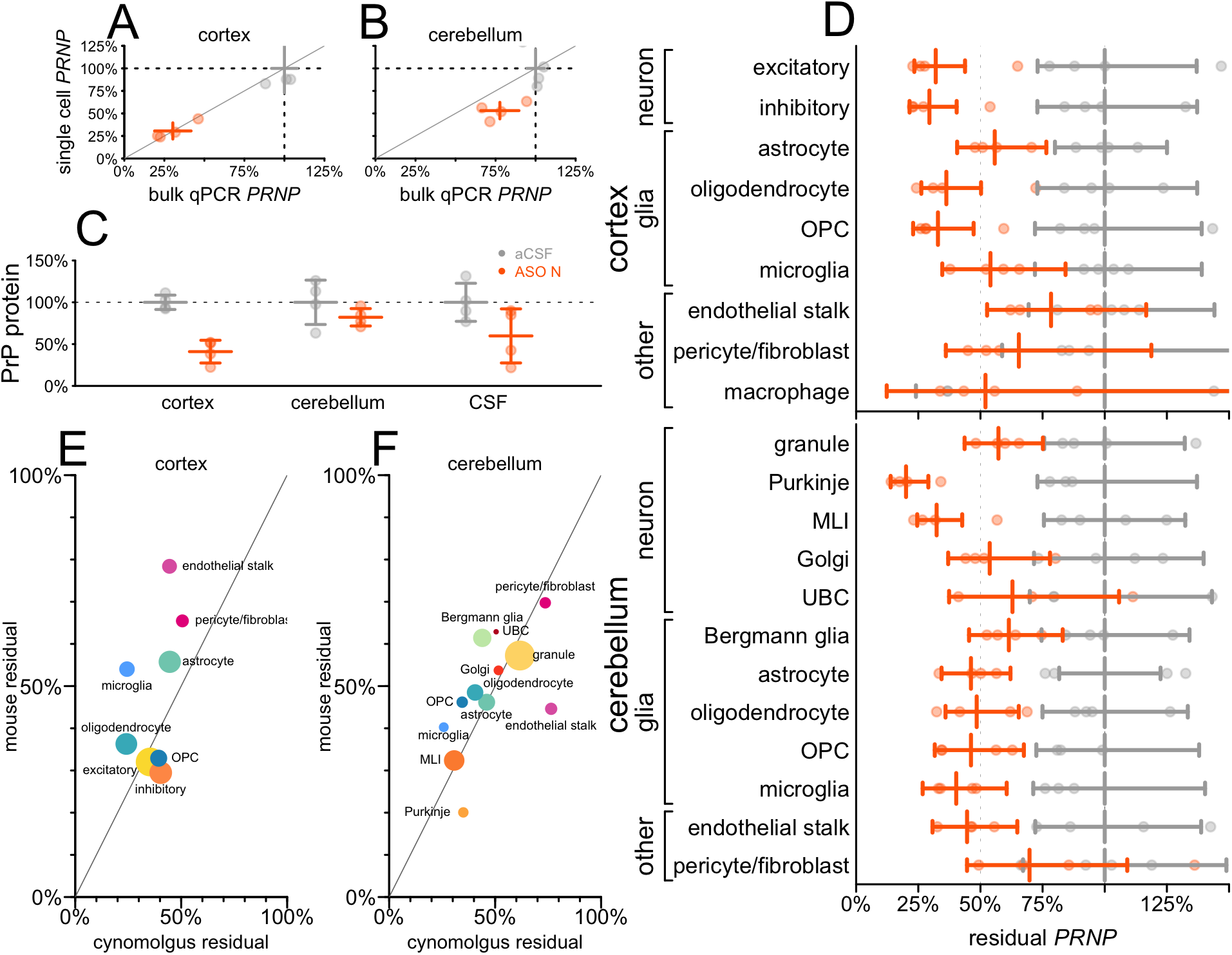
Cell type activity distribution and biomarker response in non-human primates. N = 4 cynomolgus macaques per group received 20 mg ASO N at weeks 0, 4, 8, and 12, and cortex and cerebellum were evaluated at week 13. **A-B)** PRNP knockdown assessed by bulk qPCR (x axis) versus aggregation of single cell sequencing regardless of cell type (y axis), for individual animals (points) and groups (crosshairs indicate means and 95% confidence intervals) in cortex (A) and cerebellum (B). **C)** PrP protein measured by in-house ELISA^43^ in brain parenchyma (cortex, cerebellum) and in cerebrospinal fluid (CSF). **D)** Residual PRNP by cell type in cortex and cerebellum. **E-F)** Scatterplot of cynomolgus 13-week residual PRNP by cell type (x axis) versus mouse residual Prnp at 2 weeks after a single dose of ASO 6 (data from Figure 3A) in cortex (E) and cerebellum (F). The data point for pericyte/fibroblast reflects the weighted average of residual Prnp in these two cell populations in mouse. Each point is a cell type, sized logarithmically by total number of cells (cynomolgus + mouse datasets) and colored as in Figure 1.

### Comparison of cell type specificity across paradigms

We next asked more broadly how well the profile of target engagement across cell types was shared among all of the conditions examined herein. To normalize between ASOs and timepoints with different levels of target engagement, we defined a difference from overall residual as a cell type ‘s residual target RNA, expressed as a percentage of control animals, minus the overall residual target RNA across all cell types (Table S15). Thus, positive differences indicate that a cell type has weaker knockdown than the bulk tissue, while negative differences mean deeper knockdown. In cortex, the most abundant cell types, chiefly neurons, clustered near 0% (excitatory neurons, mean +1%, inhibitory neurons, mean +2%), reflecting the bulk tissue closely, while outliers were relatively rarer cell types for which our confidence intervals on the amount of residual target are wider (Figure 6A). In cerebellum, by contrast, large differences were observed between granule cells (mean +7%) and the next two most abundant cell types, MLIs (−30%) and Bergmann glia (−10%). Overall, variability across cell types was lower in cortex and thalamus (mean weighted standard deviation 7% for both) than in cerebellum (mean weighted standard deviation 12%; Figure 6A). Accordingly, when we tested the correlation across cell types between every pair of datasets within each tissue, we observed mostly positive but non-significant correlations for cortex (Figure 6B; Table S15), whereas all correlations were strongly positive and significant in cerebellum (Figure 6C; Table S16). Because prion disease affects all neurons, we examined the estimated differences from overall residual for every neuronal subtype in every dataset (Figure 6D; Table S17). Neurons were generally either close to the bulk tissue residual (worst case, +12% difference for granule cells in *Malat1* ASO-treated mouse cerebellum at 12 weeks) or exhibited much deeper target engagement (−38% for MLIs in *Malat1* ASO-treated mouse cerebellum at 12 weeks). We did not observe any conditions in which any population of neurons exhibited dramatically weaker knockdown than the bulk tissue.

**Figure 6.**
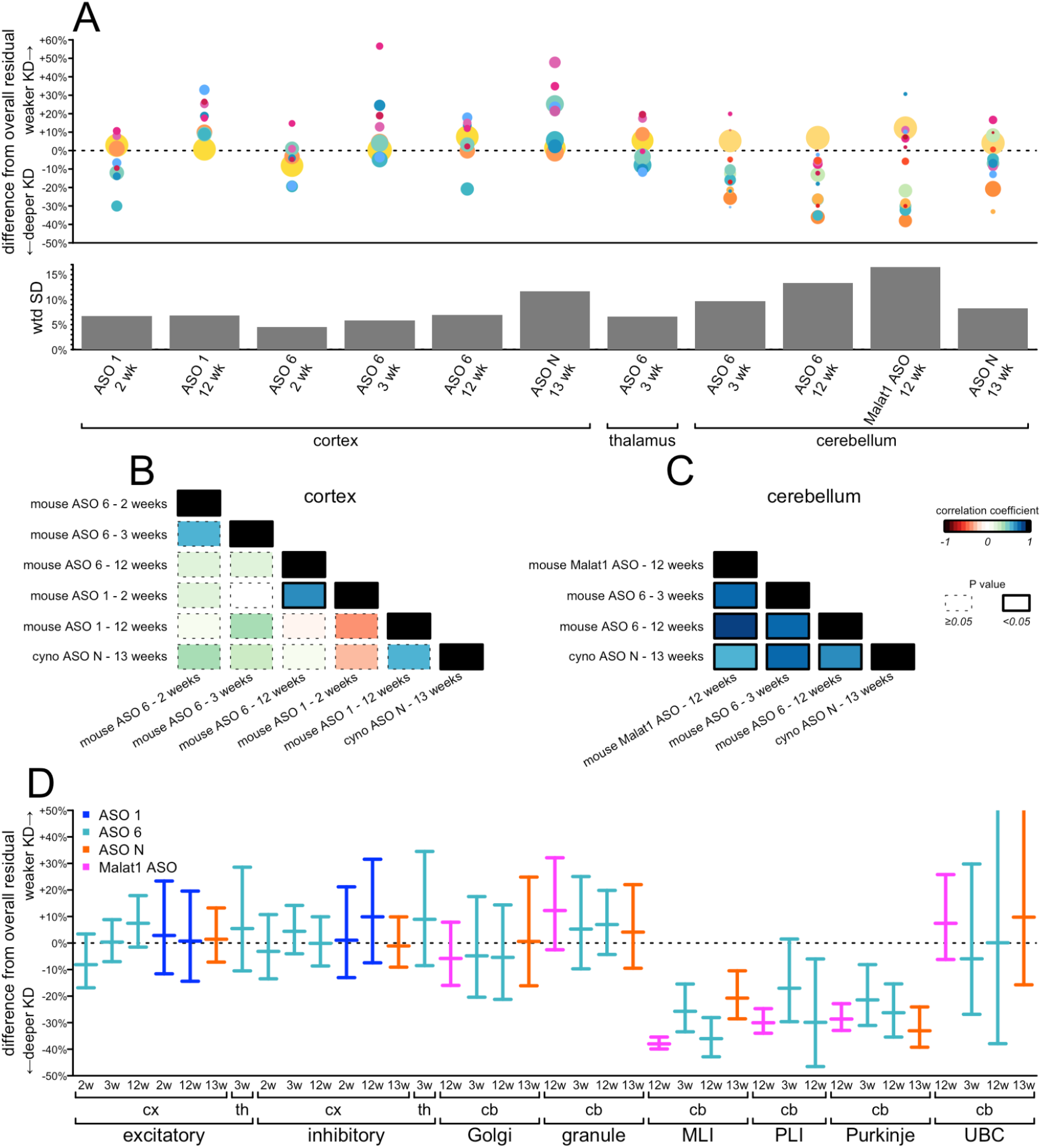
Comparison of cell type target engagement profiles across all conditions examined. **A)** Top: differences from overall residual (each cell type ‘s residual target minus the overall residual target quantified from single-nucleus data; y axis) for every combination of tissue, ASO, timepoint, and species (shared x axis). Each point is a cell type, sized logarithmically by number of cells and colored as in Figure 1. Bottom: weighted standard deviations (weighted by number of cells) in percentage points of residual target (y axis) for each condition (shared x axis). **B-C)** Correlograms (weighted Pearson ‘s correlations) between every pair of datasets for cortex (B) and cerebellum (C). Colors represent the value of the correlation coefficient rho and the outline represents the nominal P value, see legend at far right. **D)** Differences from overall residual for all neuronal subtypes in all conditions. Error bars are 95% confidence intervals from the negative binomial models for the subtype-specific residuals.

## DISCUSSION

Theoretically, 50% knockdown in bulk tissue could result, in the most extreme cases, from either 100% knockdown in half of cells or 50% knockdown in all cells. A longstanding question is where the activity of CNS ASOs falls on this spectrum. By examining the distribution of target RNA counts per cell in single nucleus sequencing data, we provide evidence of ASO activity in every single cell in a bulk tissue, albeit with differences in degree between cell types. This should be expected based on the number of drug molecules contained in a dose of ASO. For a mouse with ∼10^8^ brain cells^48,49^, a 50 µg dose of a ∼7 kDa ASO, or ∼4 ✕ 10^15^ ASO molecules, is >10^7^ molecules per brain cell. Only a small minority of ASO molecules are believed to undergo productive uptake^3^, but even if this figure is 1%, then 10^5^ productive ASO molecules per brain cell is a sufficient number that it is unlikely that any cells would avoid ASO activity simply by chance. In cynomolgus macaques, ASO concentration in some deep brain regions remains below the lower limit of quantification after intrathecal dosing^12^. Thus, in a large brain, there may be cells lacking any appreciable ASO activity due to limited drug distribution. The compounds, dosing regimes, and tissues that we chose to analyze were chosen to select scenarios with robust target engagement at the bulk tissue level, and our data argue that under this precondition, ASO activity is very broadly distributed across individual cells. This property of ASOs could prove markedly different from some gene therapy approaches to CNS diseases. In mice, engineered viral vectors for gene delivery may transduce ∼50% of CNS neurons^50^, and DNA-targeted therapeutics, with only 2 targets per cell, could provide nearly complete target suppression in those cells that are transduced. If so, modalities exhibiting similar levels of bulk target engagement could reflect rather distinct distributions at the single-cell level. These contrasting profiles might in turn present opposing challenges and opportunities for different targets.

ASOs are internalized by binding cell surface proteins^51^ and travel through different cellular uptake pathways, only a subset of which are productive, eventually escaping from endosomes to bind their targets and (for gapmer ASOs) engage RNAse H1 in the cytosol and nucleus^3^. This process presents many potential opportunities for distinct cell types to exhibit differential ASO activity, including: differences in total uptake, in the proportion of productive uptake, in the kinetics of endosomal escape, and in the rate of RNAse H1 cleavage. Histological analysis of ASO-treated brain tissue has suggested that the difference in ASO activity between granule and Purkinje cells that we observe here is likely associated with a difference in total ASO uptake^12,19^. Whether total uptake explains all of the cell type differences we observe remains to be seen.

Across cell types in the mouse cortex, deeper initial target engagement at 2 weeks appeared to correlate with more washout by 12 weeks. This correlation is expected to some degree, because target expression after washout should never recover to >100% of the untreated condition, but may also suggest that deeper initial knockdown in some cell types does not necessarily indicate a longer-lasting endosomal repository of compound.

Our dataset is ill-suited to ask genome-wide questions such as which specific cell surface proteins are most important for uptake, because any two cell types differ in the expression of many markers, not just one, and in addition, the thousands of possible answers present a large multiple testing burden which cannot be overcome by analyzing the small number of distinct cell types detected here. Cell size might be inversely related to the surface area to volume ratio, and thus to the amount of opportunity for cell surface protein binding, but UMIs/cell, a proxy for cell size^26^, was not correlated with ASO activity in our dataset. RNase H1 expression varied little across cell types and neither RNase H1 nor target expression correlated with initial target engagement or washout. In fact, this should be expected based on the number of drug molecules per cell. PrP RNA expression is on the order of hundreds of transcripts per million^52^, so a cell with 10^5^ mRNA molecules might have just tens of PrP mRNA molecules, not nearly enough to saturate 10^5^ productively uptaken ASO molecules.

Our study has many limitations. The expense of single-cell sequencing limited us to small cohort sizes (usually N = 4). For some rarer cell types, just a handful of cells per sample were observed. Many steps including nuclei dissociation, flow cytometry, and library construction, can all yield variability in number of cells and number of sequencing reads per sample. All of these factors combine to make the confidence intervals on our estimates of knockdown in many cell types rather large. Certain key observations replicate across our datasets — particularly the broadness of target engagement across cell types, with weaker knockdown in granule cells and deeper knockdown in Purkinje and interlayer neurons, and the generally weaker knockdown in cells of the vasculature. However, we studied only 3 brain regions, 4 ASOs, 2 targets, and 2 animal species, so it remains to be determined just how broadly these findings may generalize. We lack any method of quantifying drug concentration in the same cells that are sequenced, so are unable to answer questions about the pharmacokinetic/pharmacodynamic relationship at the single-cell level. Because we relied on purification of nuclei from frozen tissue, we were only able to measure target engagement in the nucleus. It is reassuring that the percentage knockdown measured by snRNA-seq was generally close to the value obtained by qPCR. However, it is not possible to perform snRNA-seq and qPCR on the exact same piece of tissue, so we instead compared adjacent pieces of tissue. On the occasions where these values diverge, we are uncertain whether it represents discordance between cytosolic and nuclear outcomes, or simply regional gradients in target engagement.

Pharmacologic interventions are seldom trialed in pre-symptomatic individuals at risk for neurodegenerative disease^53^. Observing clinical endpoints in such individuals may require lengthy follow-up^54^ or may be outright numerically infeasible^55^. This has led to the suggestion that in prion disease, where the central role of PrP in disease is incontrovertible^56^, the lowering of CSF PrP — a target engagement biomarker only — could serve as a primary endpoint in trials of at-risk individuals^21^. This prospect demands that especially strong data from animal studies will be needed to certify the links between CSF PrP, target engagement in the disease-relevant cells, and modification of disease^57^. In prion disease, the critical cells to engage are neurons. Although astrocytes may contribute to disease by propagating prions^58–60^, only neurons degenerate in prion disease, and neurotoxicity is cell autonomous: neurons that do not express PrP are protected even if they are in direct contact with misfolded prions produced by neighboring cells^60–62^. In contrast, neuroinflammatory responses from astrocytes and microglia^63–66^ appear to be strictly non-autonomous, requiring neuronal prion infection^60^. That PrP-lowering ASOs extend survival in prion-infected mice^22,41,67^ implies that they must lower PrP in neurons; nevertheless, we felt it prudent to further examine this link to determine whether there might ever exist circumstances in which a bulk tissue readout would indicate PrP lowering despite little or no target engagement in neurons. It is reassuring, then, that across a range of experimental parameters — dosing regimens, times post-dose, ASO chemistries and gapmer configurations, targets, species, and brain regions — we never identified a circumstance in which bulk tissue would misinform about PrP RNA having been lowered in neurons. Of course, given the somewhat differing activity of ASOs in distinct CNS cell types and the potential for drug concentration gradients across the brain, no single compartment readout, such as CSF PrP, can accurately report on every disease-relevant cell in this whole brain disease. Still, our findings provide one pillar of support for the expectation that lowered CSF PrP in an ASO trial is reasonably likely to predict clinical benefit in individuals at risk for prion disease.

## ACKNOWLEDGMENTS

This study was funded primarily by the Ono Pharma Foundation (Oligonucleotide Medicine Award 2019) and by Ionis Pharmaceuticals. The authors also acknowledge support from the National Institutes of Health (R21 TR003040 and R01 NS125255). The authors are grateful to Bo Li, Brian Granger, Brittany Ford, Briana Buscemi, and the Broad Institute Flow Cytometry Facility for technical assistance, to Joe Schroeder and Tom Zanardi for monitoring the NHP study, and to C. Frank Bennett for critical feedback on the manuscript.

## SUPPLEMENTARY MATERIAL

Tables S1 - S17 are available in the attached Excel file or as tab-separated text files at https://github.com/ericminikel/scaso

**Figure S1.**
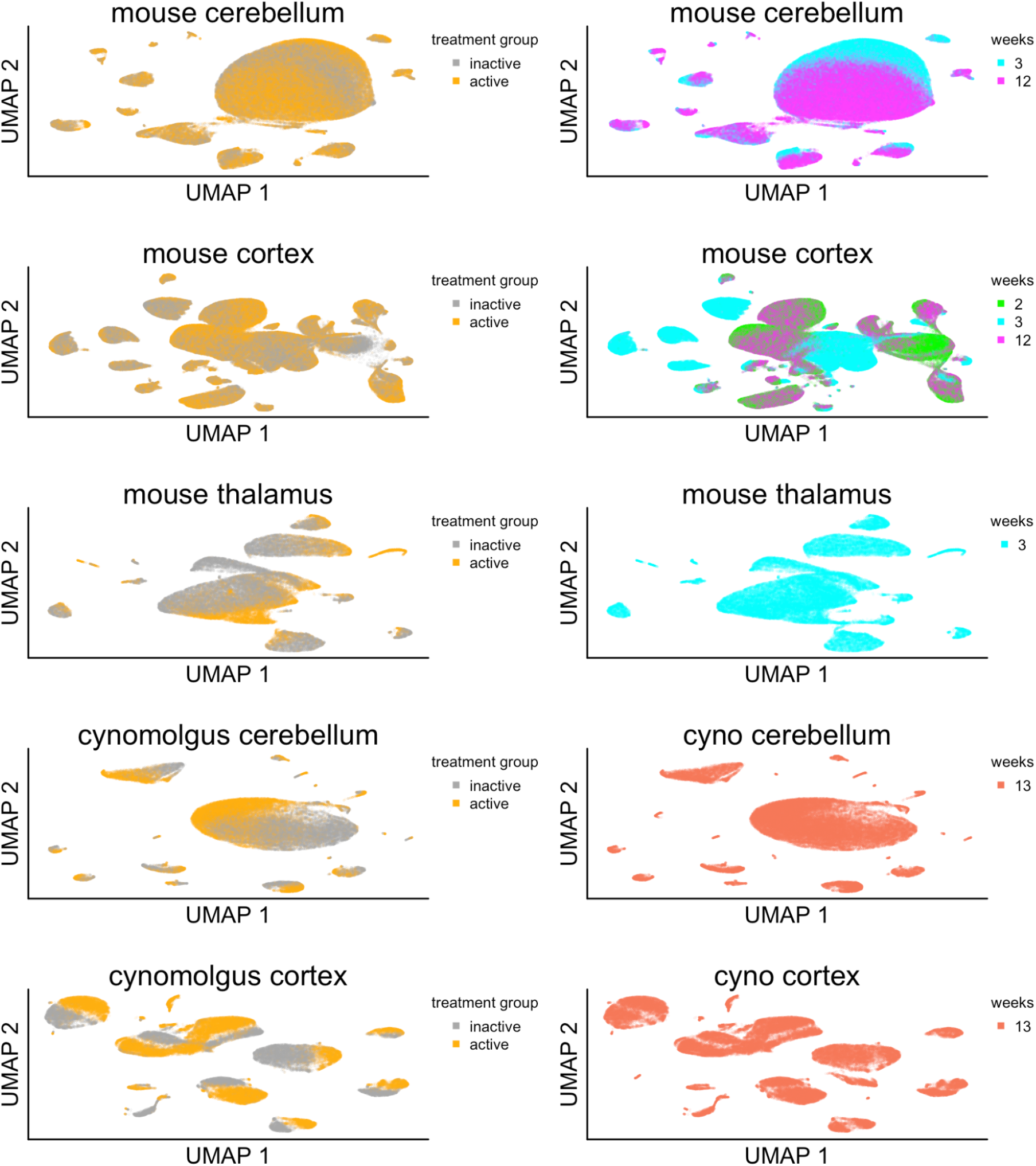
Weeks post-dose and treatment group in UMAP space. UMAP plots from Figure 1 colored by treatment group (left) or weeks post-dose (right). Twinning of some cell types, particularly in mouse cortex, is due to a batch effect between 3-week versus 2- and 12-week post-dose animals; clusters are generally well-balanced between active and inactive treatment groups.

